# Distinct antagonist-bound inactive states underlie the divergence in the structures of the dopamine D2 and D3 receptors

**DOI:** 10.1101/640870

**Authors:** J. Robert Lane, Ara M. Abramyan, Ravi Kumar Verma, Herman D. Lim, Jonathan A. Javitch, Lei Shi

## Abstract

Understanding how crystal structures reflect the range of possible G protein-coupled receptor (GPCR) states is critical for rational drug discovery (RDD). Combining computational simulations with mutagenesis and binding studies, we find that the structure of the dopamine D2 receptor (D_2_R)/risperidone complex captures an inactive receptor conformation that accommodates some but not all antagonist scaffolds. Indeed, we find that eticlopride binds D_2_R in a configuration very similar to that seen in the D_3_R structure, in a pose that is incompatible with the D_2_R/risperidone structure. Moreover, our simulations reveal that extracellular loops 1 and 2 (EL1 and EL2) are highly dynamic, with spontaneous transitions of EL2 from the helical conformation in the D_2_R/risperidone structure to an extended conformation similar to that in the D_3_R/eticlopride structure. Our results highlight previously unappreciated conformational diversity and dynamics in the inactive state of a GPCR with potential functional implications. These findings are also of paramount importance for RDD as limiting a virtual screen to one state will miss relevant ligands.

## INTRODUCTION

G protein-coupled receptors (GPCRs) are important therapeutic targets for numerous human diseases. Our understanding of GPCR functional mechanisms has evolved from a simple demarcation of single active and inactive states to the appreciation and detection of multiple active states responsible for partial or biased agonism^1–3^. High-resolution crystal structures of these proteins are vital for structure-based (rational) drug discovery (RDD) efforts designed to tailor selectivity and efficacy^4,5^. Indeed, a current focus of the field is on developing functionally biased ligands^6–8^ that couple preferentially to a particular effector pathway. Less attention has been dedicated to the possibility that there may be multiple inactive states, and that different antagonist scaffolds might lead to different receptor conformations. Such a possibility could have major impact on RDD for antagonists, since a crystal structure of a receptor with a particular ligand bound might represent an invalid docking target for an antagonist that binds in a different pose to a different inactive conformation. Although substantial differences in antagonist binding mode and position of the binding pockets have been revealed between different aminergic receptors, no differences in conformation has been detected for the inactive state of any individual aminergic receptor^5^. In particular, although a number of antagonists from different scaffolds have been co-crystallized with β_2_ adrenergic receptor, the conformational differences among these crystal structures are minimal^5^.

Curiously, the structures of the highly homologous dopamine D2 and D3 receptors (D_2_R and D_3_R) in the inactive states revealed quite substantial differences on the extracellular side of the transmembrane domain when bound with antagonists from different scaffolds^9,10^. Specifically, the D_3_R structure is in complex with eticlopride, a substituted benzamide (PDB: 3PBL)^9^, while the D_2_R structure is bound with risperidone, a benzisoxazole derivative (PDB: 6CM4)^10^. The binding poses of the two ligands differ substantially. The risperidone is oriented relatively perpendicular to the membrane plane with its benzisoxazole ring penetrating into a hydrophobic pocket beneath the orthosteric binding site (OBS) of D_2_R; in contrast, eticlopride is oriented relatively parallel to the membrane plane contacting the extracellular portion of TM5 in D_3_R that risperidone does not touch in D_2_R^10,11^. Nemonapride, another substituted benzamide, binds in the OBS of the slightly divergent D_4_R (PDB: 5WIV)^12^ in a manner very similar to that of eticlopride in the D_3_R^11^. The benzisoxazole moiety of risperidone is enclosed by 8 residues in D_2_R that are identical among all D2-like receptors (i.e., D_2_R, D_3_R, and D_4_R): Cys118^3.36^ (superscripts denote Ballesteros-Weinstein numbering^13^), Thr119^3.37^, Ile122^3.40^, Ser197^5.46^, Phe198^5.47^, Phe382^6.44^, Trp386^6.48^, and Phe390^6.52^. Notably, three of these residues on the intracellular side of the OBS, Ile122^3.40^, Phe198^5.47^, Phe382^6.44^, accommodate the F-substitution at the tip of the benzisoxazole ring in a small cavity (termed herein as the Ile^3.40^ sub-pocket) (Fig. 1a). Interestingly, this Ile^3.40^ sub-pocket is collapsed in both the D_3_R and D_4_R structures^11^ (Fig. 1b,c). We noted that this collapse is associated with rotation of the sidechain of Cys^3.36^: In the D_2_R/risperidone structure, the sidechain of Cys^3.36^ faces the OBS, whereas it rotates downwards to partially fill the Ile^3.40^ sub-pocket in the D_3_R/eticlopride and D_4_R/nemonapride structures.

**Figure 1.**
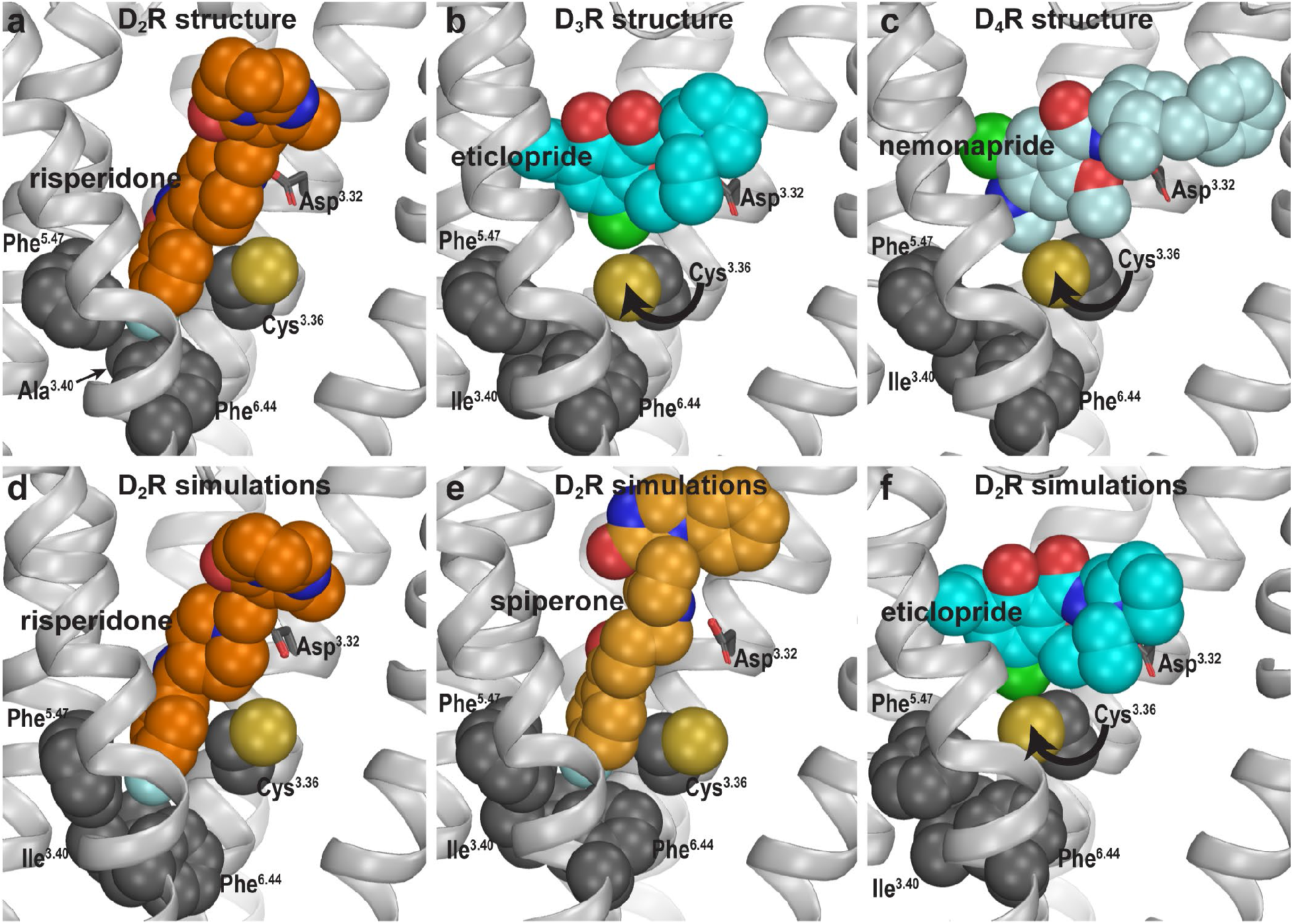
Divergent occupations of the Ile^3.40^ sub-pocket by non-selective ligands from different scaffolds. In the D_2_R structure (**a**), the F-substitution on the benzisoxazole ring of risperidone occupies the Ile^3.40^ sub-pocket enclosed by conserved Ile^3.40^, Phe^5.47^, and Phe^6.44^. The same viewing angle shows that in the D_3_R (**b**) and D_4_R (**c**) structures, Cys^3.36^ rotates to fill in the Ile^3.40^ sub-pocket, and the substituted benzamides eticlopride and nemonapride cannot occupy the aligned sub-pockets. In our D_2_R/risperidone simulations (**d**), risperidone maintains its pose revealed by the crystal structure. In the D_2_R/spiperone simulations (**e**), the Ile^3.40^ sub-pocket is similarly occupied as in D_2_R/risperidone. In the D_2_R/eticlopride simulations (**f**), the Ile^3.40^ sub-pocket is collapsed as in the D_3_R (**b**) and D_4_R (**c**) structures (this trend is independent of the force field being used in the simulations).

Importantly, the co-crystalized ligands (risperidone, eticlopride, and nemonapride) display little subtype selectivity across D_2_-like receptors^9,12,14,15^ (also see PDSP database^16^). Given the high homology among D_2_-like receptors, especially between D_2_R and D_3_R in and near the OBS, we hypothesized that the drastic conformational differences between the structures of these receptors in the inactive state are mostly due to the different binding poses of antagonists of different scaffolds and not to inherent differences between the two receptors. To test this hypothesis, we carried out extensive molecular dynamics (MD) simulations of D_2_R in complex with non-selective antagonists of different scaffolds to characterize the plasticity of the OBS and the extracellular loop dynamics in the inactive conformational state.

## RESULTS

In the D_2_R structure, Ile122^3.40^, Leu375^6.37^, and Leu379^6.41^ were mutated to Ala to thermostabilize the receptor for crystallography. We reverted these thermostabilizing mutations back to their WT residues and established WT D_2_R models in complex with selected ligands (see Methods, Supplementary Fig. 1 and Supplementary Table 1).

### Spiperone but not eticlopride extends into the Ile^3.40^ sub-pocket

In our MD simulations of the WT D_2_R/risperidone complex, we observed that risperidone stably maintains the binding pose captured in the crystal structure, even without the thermostabilizing mutations (Fig. 1d). Thus, the I122^3.40^A mutation has minimal impact on the binding pose of risperidone. Interestingly in the simulations of the D_2_R model in complex with spiperone, a butyrophenone derivative, the F-substitution on the butyrophenone ring similarly occupies the Ile^3.40^ sub-pocket as risperidone (Fig. 1e). Note that the F-substitutions in risperidone and spiperone are located at similar distances to the protonated N atoms that interact with Asp^3.32^ (measured by the number of carbon atoms between them, Supplementary Fig. 1) and these two ligands appear to be optimized to occupy the Ile^3.40^ sub-pocket.

In contrast, in our simulations of the D_2_R/eticlopride complex, the eticlopride pose revealed in the D_3_R structure (PDB: 3PBL) is stable throughout the simulations and does not protrude into the Ile^3.40^ sub-pocket (Fig. 1f). Consistent with the difference in the crystal structures noted above (Fig. 1a,b), when risperidone and spiperone occupy the Ile^3.40^ sub-pocket, the sidechain of Cys118^3.36^ rotates away with its χ1 rotamer in *gauche-*, while in the presence of the bound eticlopride, this rotamer is in *trans* (Supplementary Fig. 2).

### Mutagenesis of Ile122^3.40^ and the sensitivity to Na^+^

To validate these computational findings, we mutated Ile122^3.40^ of WT D_2_R to both Trp and Ala and characterized how these mutations would affect the binding affinities for spiperone, risperidone, and eticlopride (Table 1). We hypothesized that the bulkier sidechain of Trp at position 3.40 would hamper the binding of spiperone and risperidone since they occupy the Ile^3.40^ sub-pocket but have no effect on eticlopride binding, while the smaller Ala should not affect the binding of spiperone or risperidone. Consistent with this hypothesis, compared to WT, the I122W mutation significantly decreased the binding affinities of risperidone and spiperone but had no effect on that of eticlopride. In contrast, the I122A mutation did not affect the affinities of spiperone or risperidone but caused a 3-fold increase in the affinity of eticlopride. This is consistent with our simulation results that show the I122^3.40^A mutation has minimal impact on risperidone binding. Both Ile122^3.40^ and Phe382^6.44^ of the Ile122^3.40^ sub-pocket are part of the conserved Pro^5.50^-Ile^3.40^-Phe^6.44^ motif that undergoes rearrangement upon receptor activation^17^, and we have found that the I122A mutation renders the receptor non-functional^10,18^. Thus, the I122A mutation may promote an inactive conformation of D_2_R that favors eticlopride binding but has no effect on the binding of risperidone or spiperone. This is consistent with our proposal that different antagonist scaffolds may favor distinct inactive conformations of D_2_R.

**Table 1.** The effect of mutations on the binding affinities of selected D_2_R ligands. The affinities of [^3^H]spiperone were determined in saturation experiments at WT or mutant SNAP-tagged D2SRs stably expressed in FlpIN CHO cells. Binding affinity values for aripiprazole and risperidone were obtained in competition binding experiments. Values are expressed as mean ± S.E.M. from three separate experiments. *P<0.05, significantly different from the wild-type receptor determined by a one-way ANOVA, Dunnett post-hoc test.

| SNAP-D<sub>2</sub>SR | [<sup>3</sup>H]spiperone saturation binding |  | [<sup>3</sup>H]spiperone competition binding |  |
| --- | --- | --- | --- | --- |
|  | *pK<sub>d</sub>*<br>(*K<sub>d</sub>*, nM) | *B<sub>max</sub>*<br>(pmol.mg<sup>-1</sup>) | Risperidone<br>*pK<sub>i</sub>* (*K<sub>i</sub>*, nM) | Eticlopride<br>*pK<sub>i</sub>* (*K<sub>i</sub>*, nM) |
| WT | 9.72 ± 0.11<br>(0.19) | 7.95 ± 1.63 | 8.55 ± 0.20 (2.8) | 9.84 ± 0.17<br>(0.14) |
| WT -Na<sup>+</sup> | 9.70 ± 0.11<br>(0.20) | 6.39 ± 1.04 | 8.96 ± 0.04 (1.1) | - |
| I122<sup>3.40</sup>A | 9.74 ± 0.15<br>(0.18) | 4.37 ± 0.92 | 8.10 ± 0.03 (7.9) | 10.33 ± 0.04*<br>(0.04) |
| I122<sup>3.40</sup>W | 8.95 ± 0.08*<br>(1.15) | 2.61 ± 0.50 | 7.43 ± 0.12* (37) | 9.61 ± 0.0<br>(0.25) |

We have previously shown that the binding of structurally distinct antagonists to the OBS is differentially modulated by the Na^+^ bound in a conserved allosteric binding pocket coordinated by Asp^2.50^ and Ser^3.39^. While spiperone binding was insensitive to the presence of Na^+^, the affinities of eticlopride and sulpiride were increased and the affinity of zotepine was decreased^19^. In our current simulations, the occupancy of the Ile^3.40^ sub-pocket by both spiperone and risperidone was unaffected by the presence or absence of bound Na^+^ (Supplementary Fig. 2). Note that Ser^3.39^ and Ile^3.40^ are adjacent to each other, and we hypothesized that the occupation of the Ile^3.40^ sub-pocket by spiperone or risperidone may confer a lack of Na^+^ sensitivity to the binding of these inverse agonists. To further test this hypothesis and to understand how Na^+^ influences risperidone binding, we performed binding experiments in the absence or presence of Na^+^ and found its affinity to be unaffected (Table 1).

Interestingly in our simulations, while the eticlopride pose is highly stable in the presence of bound Na^+^, eticlopride oscillated between subtly different poses in the absence of Na^+^. These oscillations are associated with the sidechain of Cys^3.36^ swinging back and forth between two rotamers, suggesting an important role of Na^+^ binding in stabilizing the eticlopride pose and the configuration of the Ile^3.40^ sub-pocket (Supplementary Fig. 2). Intriguingly, in the recently reported crystal structure of the serotonin 2A receptor (5-HT_2A_R) in complex with zotepine (PDB: 6A94)^20^, the benzothiepin moiety of zotepine also occupies the Ile^3.40^ sub-pocket, but this results in noticeable differences in the nearby residues compared to the 5-HT_2A_R/risperidone structure (PDB: 6A93)^20^. Specifically, the sidechain of the thermostabilizing mutation S162^3.39^K, which occupies the Na^+^ binding site, moves further away from Ile163^3.40^ in the presence of the bound zotepine compared to that in the presence of risperidone. Such a difference indicates that zotepine and risperidone may have different sensitivities of Na^+^ binding, which is consistent with the insensitivity of risperidone binding to Na^+^ and the higher binding affinity of zotepine in the absence of Na^+^ that we have previously observed at D_2_R^19^.

Together these findings along with our previously published data are consistent with our hypothesis that the binding of the ligands with different scaffolds stabilize distinct inactive conformations of D_2_R such that they are differentially sensitive to the presence of Na^+^, and there is likely an allosteric connection between the Na^+^ binding site and the Ile^3.40^ sub-pocket.

### Plasticity of the ligand binding site in response to ligands with different scaffolds

On the extracellular side of the OBS, the space near Ser^5.42^ and Ser^5.43^ that accommodates the bulky substitutions of the benzamide rings of bound eticlopride and nemonapride in the D_3_R and D_4_R structures is not occupied by risperidone in D_2_R. Furthermore, risperidone positions the aromatic cluster of TM6 and TM7 (Trp^6.48^, Phe^6.51^, Phe^6.52^, His^6.55^, and Tyr^7.35^) in D_2_R differently from its configurations in the D_3_R and D_4_R structures. These differences are likely associated with the inward and outward movements, respectively, of the extracellular portions of TM5 and TM6 in D_2_R relative to those in the D_3_R and D_4_R structures (Fig. 2a). To evaluate whether these conformational rearrangements are due to the minor divergence in these regions of the receptors or to the ligand binding site plasticity to accommodate ligands from different scaffolds, we compared the resulting conformations of D_2_R bound with risperidone or eticlopride. We observed the same trend of rearrangements of the transmembrane segments surrounding the OBS in the resulting receptor conformations from our D_2_R/risperidone and D_2_R/eticlopride simulations (Fig. 2b), i.e., an inward movement of TM6 and outward movement of TM5 in the presence of bound eticlopride (Fig. 2c,d). Without such movements in D_2_R/eticlopride, Ser193^5.42^ and Ser194^5.43^ would clash with the bound eticlopride (Fig. 2b). These findings further support our inference that these differences between the D_2_R and D_3_R inactive structures are largely due to the different scaffolds of the non-selective ligands bound.

**Figure 2.**
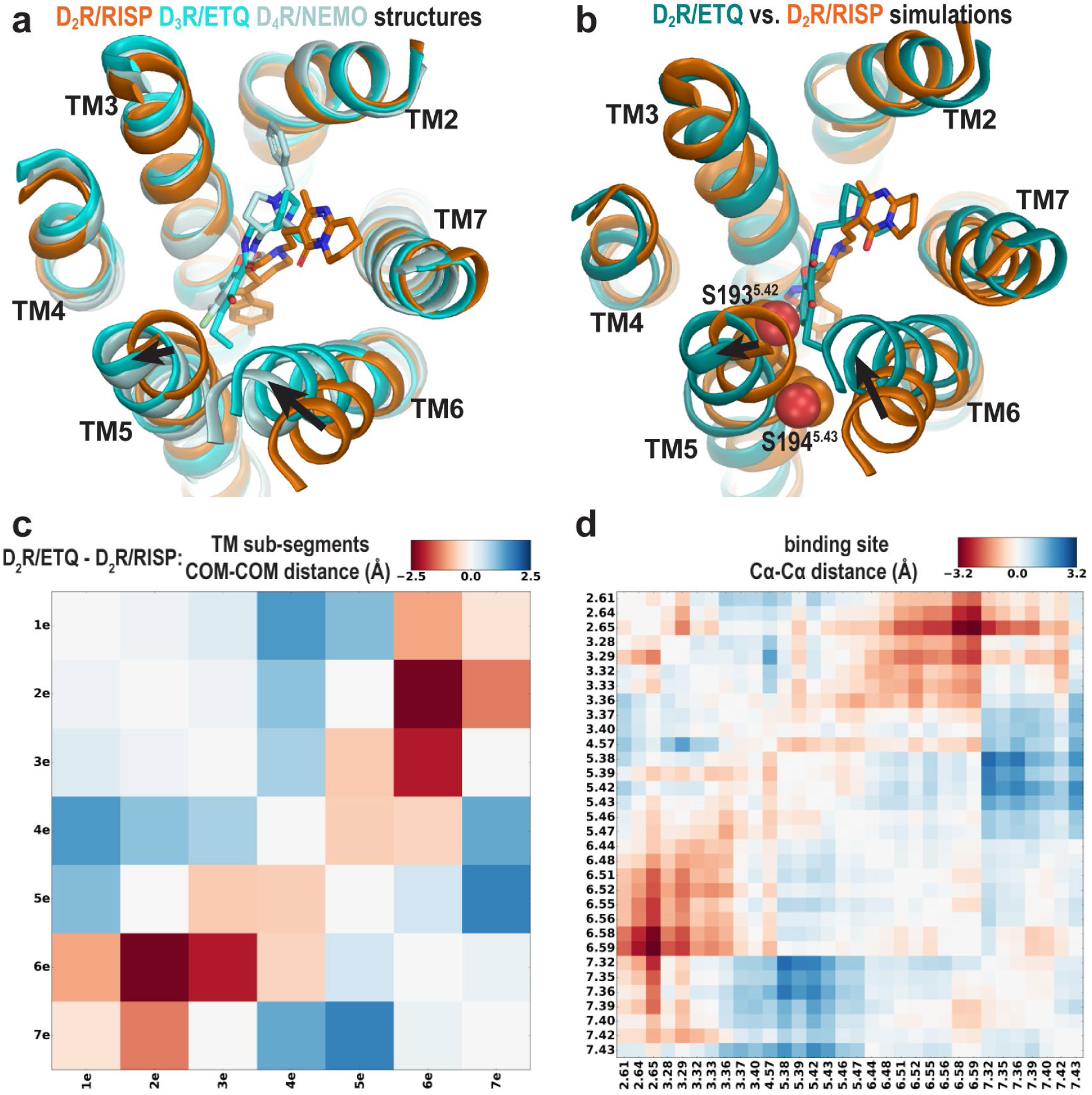
The different conformations in the extracellular vestibules of D_2_R, D_3_R, and D_4_R are likely due to binding of non-selective ligands from different scaffolds. (**a**) Superpositioning of D_2_R, D_3_R, and D_4_R structures shows that the binding of eticlopride (ETQ, cyan) in D_3_R and nemonapride (NEMO, pale cyan) in D_4_R result in outward and inward rearrangements of the extracellular portions of TM5 and TM6, respectively, compared to the binding of risperidone (RISP, orange) in D_2_R. (**b**) superpositioning of representative frames of the D_2_R/ETQ and D_2_R/RISP simulations shows a similarly trend of the outward and inward movements of TM5 and TM6, respectively, in the presence of the bound ETQ, even when the simulations were started from the D_2_R conformation stabilized by RISP. Note Ser193^5.42^ and Ser194^5.43^ would clash with the bound eticlopride if there was conformational adjustment. (**c**, **d**) PIA-GPCR analysis (see Methods) comparing the D_2_R/ETQ and D_2_R/RISP conformations. The analysis of the pairwise-distance differences among the subsegments (**c**) indicates that TM6e moves inward (smaller distance to TM2e, dark red pixel), while TM5e moves outward (larger distances to TM7e, dark blue pixel) in the D_2_R/ETQ simulations. The analysis of pairwise-distance differences among the Cα atoms of the ligand binding residues (**d**) indicates significant changes near residues Phe189^5.38^, Ser193^5.42^, Asn367^6.58^, and Ile368^6.59^ (darker colored pixels).

### The extracellular loop 2 (EL2) of D_2_R/risperidone can spontaneously unwind

In addition to these differences in the transmembrane segments surrounding the OBS, there are also substantial differences in the configuration of EL2 in the D_2_R and D_3_R structures. EL2 (residues 173 to 186) between TM4 and TM5 is connected to TM3 via a disulfide bond formed between Cys182^EL2.50^ (see Methods and Supplementary Fig. 3 for the indices of EL1 and EL2 residues) and Cys107^3.25^. The conformation of EL2, the sequence of which is not conserved among aminergic GPCRs, is expected to be dynamic. Indeed, in the D_2_R/risperidone structure, the sidechains of residues 176^EL2.40^, 178^EL2.46^, 179^EL2.47^, and 180^EL2.48^, which are distal to the OBS were not solved, likely due to their dynamic nature. Curiously the portion of EL2 C-terminal to Cys182^EL2.50^ (residues 182^EL2.50^-186^EL2.54^), which forms the upper portion of the OBS that is in contact with ligand, is in a helical conformation in the D_2_R/risperidone structure.

Strikingly, in our MD simulations, we found that this helical region showed a tendency to unwind (Supplementary Movie 1). The unwinding of EL2 involves a drastic rearrangement of the sidechain of Ile183^EL2.51^, which must dissociate from a hydrophobic pocket formed by the sidechains of Val111^3.29^, Leu170^4.60^, Leu174^EL2.38^, and Phe189^5.38^. This process is initiated by the loss of a hydrogen-bond (H-bond) interaction between the sidechain of Asp108^3.26^ and the backbone amine group of Ile183^EL2.51^ formed in the D_2_R/risperidone structure (Fig. 3b, step i). When this interaction is broken, the orientation of residues 182^EL2.50^-186^EL2.54^ deviates markedly from that of the crystal structure, losing its helical conformation (see below). Subsequently the sidechain of Ile183^EL2.51^ rotates outwards and passes a small steric barrier of Gly173^EL2.37^ (Fig. 3b, step ii), and in some trajectories makes a favorable hydrophobic interaction with the sidechain of Ala177^EL2.45^. In a few long trajectories, Ile183^EL2.51^ rotates further towards the extracellular vestibule where it can make favorable interactions with hydrophobic or aromatic residues from the N terminus, or the bound risperidone (Supplementary Movie 1). Consequently, residues 182^EL2.50^-186^EL2.54^ are in a fully extended loop conformation while Ile184^EL2.52^ tilts under EL2 (Fig. 3b, step iii).

**Figure 3.**
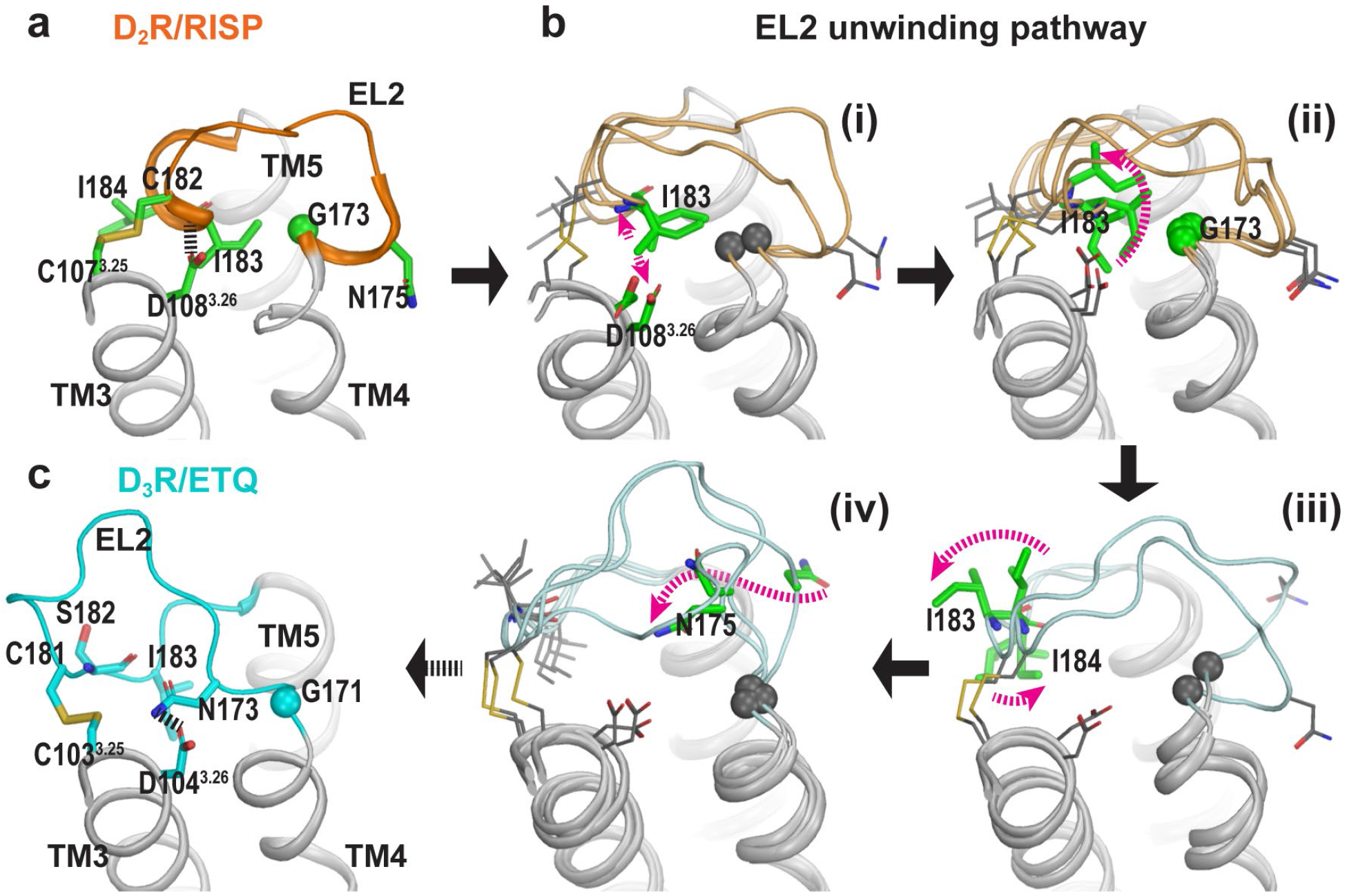
The helical region of EL2 of D_2_R can spontaneously unwind to an extended conformation similar to that of D_3_R. (**a**) Residues 182^EL2.50^-186^EL2.54^ in the D_2_R/RISP structure are in a helical conformation. EL2 is connected to TM3 via a disulfide bond (Cys182^EL2.50^-Cys107^3.25^), while the backbone of Ile183^EL2.51^ forms an interaction with Asp108^3.26^ (magenta dotted line). (**b**) The key events in the EL2 unwinding pathway (for each step, a number of representative frames are shown): the ionic interaction between Asp108^3.26^ and Ile183^EL2.51^ has to dissociate first (**i**), which allows the sidechain of Ile183 to rotate towards lipids and pass through a minor barrier formed by Gly173^EL2.37^ (**ii**); then the sidechain of Ile183^EL2.51^ rotates towards the extracellular vestibule while that of Ile184^EL2.52^ tilts under EL2 (**iii**); these changes allow Asn175^EL2.39^ to move from facing lipid to facing the binding site (**iv**). The resulting conformation of EL2 of D_2_R is similar to that of D_3_R for all the aforementioned residues (**c**). In particular, Asn173^EL2.39^ of D_3_R, which aligns to Asn175^EL2.39^ of D_2_R, forms an H-bond interaction with Asp104^3.26^.

In the D_3_R structure, the aligned residue for Asp108^3.26^ of D_2_R is conserved as Asp104^3.26^; its sidechain forms an interaction not with Ile182^EL2.51^ but rather with the sidechain of Asn173^EL2.39^, which is also conserved in D_2_R as Asn175^EL2.39^. In the D_4_R, the aligned two residues (Asp109^3.26^ and Asn175^EL2.39^) are conserved as well, their sidechains are only 4.3 Å away in the D_4_R structure, slightly larger than 3.2 Å in the D_3_R structure. Even though these residues are conserved in D_2_R, the interaction in D_3_R (and potentially in D_4_R), between Asp^3.26^-Asn^EL2.39^, is not present in the D_2_R structure in which the aligned Asn175^EL2.39^ faces lipid (Fig. 3a). However, in a few of our long D_2_R simulations, Asn175^EL2.39^ gradually moves inwards and approaches Asp108^3.26^ (Fig. 3b, step iv). At this point, the EL2 conformation of D_2_R is highly similar to that of D_3_R (Fig. 3c), suggesting that EL2 is dynamic and can exist in both conformations.

### Both EL2 conformation and ligand scaffold affect the EL1 conformation

We have previously shown that the divergence in both the length and number of charged residues in EL1 among D_2_R, D_3_R, and D_4_R is responsible for the selectivity of more extended ligands^21,22^. Another striking difference in the D_2_R, D_3_R, and D_4_R structures is the position of the conserved Trp^EL1.50^ in EL1. Trp100^EL1.50^ is in a much more inward position in the D_2_R structure, making a direct contact with the bound risperidone (Fig. 4a), Trp101^EL1.50^ in D_4_R interacts with the bound nemonapride that has an extended structure, whereas Trp96^EL1.50^ in D_3_R is not in contact with eticlopride (Fig. 4b). Thus, we asked whether these distinct positions of Trp^EL1.50^ are due to the divergence in EL1 among these receptors^21^ or due to the multiple inactive conformations that accommodate the binding of non-selective ligands of divergent scaffolds.

**Figure 4.**
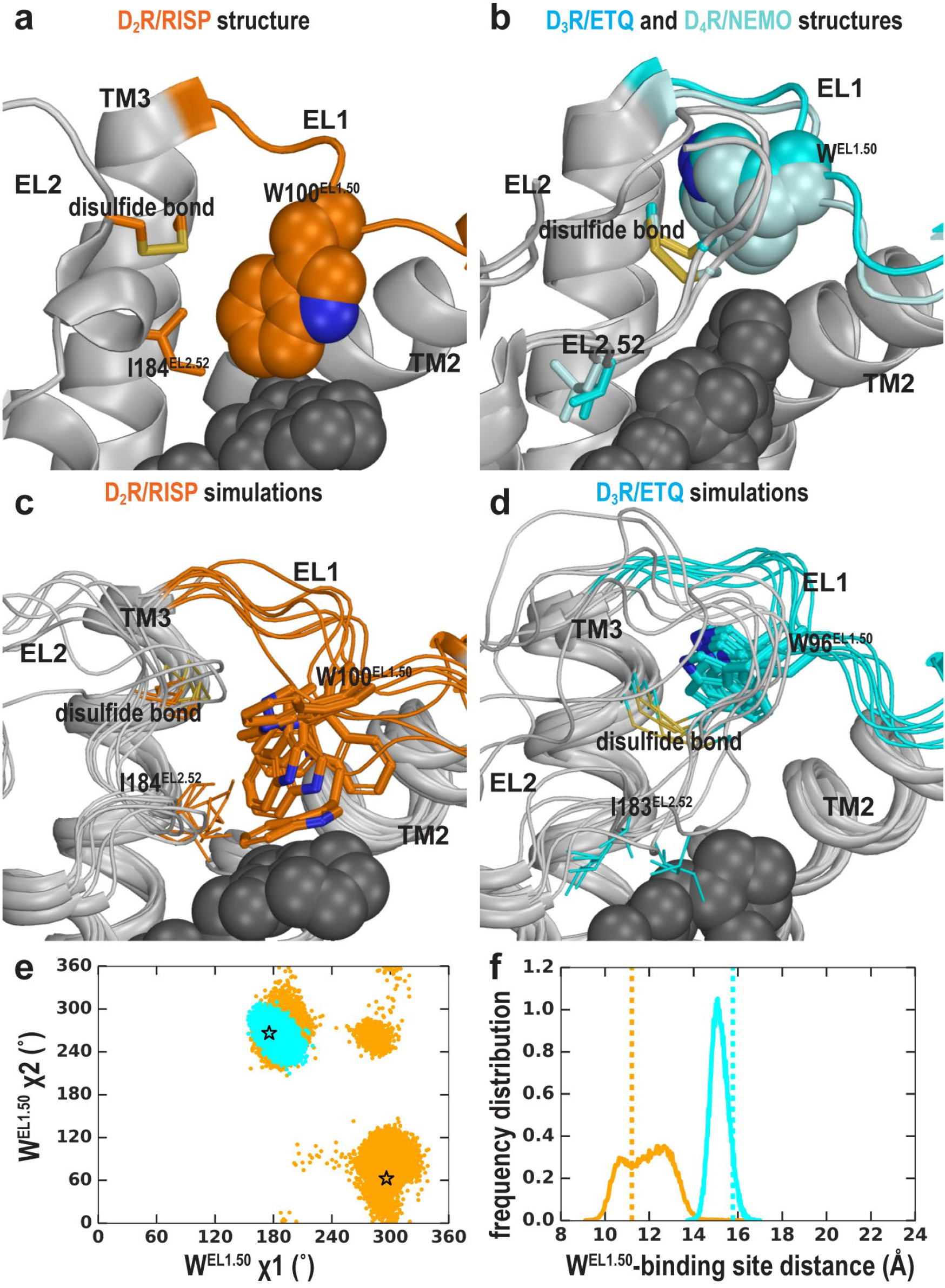
The EL2 conformation affects the EL1 conformation. Divergent EL1-EL2 interfaces among the D_2_R (**a**), D_3_R, and D_4_R (**b**) structures. In the D_2_R structure, the Trp100^EL1.50^ in EL1 only forms a weak interaction with Ile184^EL1.52^; while the aligned Trp96^EL1.50^ of D_3_R and Trp101^EL1.50^ in D_4_R are stabilized by their interactions with the disulfide bond – their passages towards the position of Trp100^EL1.50^ in D_2_R are blocked by the extended EL2. In our simulations, Trp100^EL1.50^ in D_2_R shows significant flexibility and can adopt multiple positions and orientations in D_2_R/risperidone (**c**), while Trp96^EL1.50^ in D_3_R is highly stable in D_3_R/eticlopride (**d**). (**e**) The χ1 and χ2 dihedral angles of Trp100^EL1.50^ in the subset of the D_2_R/risperidone simulations in which EL2 is still in a helical conformation (orange), are more widely distributed than those of Trp96^EL1.50^ in the D_3_R/eticlopride simulations in which EL2 remains in extended conformations (cyan). These dihedral angle values in the D_2_R and D_3_R structures are indicated with the orange and cyan stars, respectively. (**f**), For the same two sets of simulations in **e**, the distance between the center of mass (COM) of the sidechain heavy atoms of Trp100 in D_2_R and the COM of the Cα atoms of the ligand binding site residues (excluding Trp100, see Methods for the list of the residues) has wider distributions than the corresponding distance between Trp96^EL1.50^ in D_3_R and its ligand binding site. These distances in the D_2_R and D_3_R structures are indicated with the orange and cyan dotted lines, respectively.

In the D_2_R/risperidone simulations, we found that when residues 182^EL2.50^-186^EL2.54^ of EL2 are in a helical conformation and there is more room in the extracellular vestibule, the position of Trp100^EL1.50^ is flexible and can adopt several positions and orientations (Fig. 4c,e,f). In the D_2_R/eticlopride simulations, when EL2 is helical, Trp100^EL1.50^, which cannot interact with eticlopride, shows more flexibility and can move to a similar position like that of Trp96^EL1.50^ in the D_3_R structure (Supplementary Fig. 4 and Supplementary Movie 2). Interestingly, in this position, the conformation of Trp^EL1.50^ can be stabilized by the disulfide bond of EL2^23^ as shown in Supplementary Movie 2 or by interaction with the N terminus, which was truncated in the receptor used for the crystal structure. In the D_2_R/spiperone simulations, the phenyl substitution on the triazaspiro[4.5]decane moiety protrudes towards the interface between TM2 and TM3, and contacts Trp100^EL1.50^, which is flexible as well but can adopt a position that is even further away from the OBS than that of Trp96^EL1.50^ in the D_3_R structure (Supplementary Fig. 4).

In contrast, when EL2 is in an extended conformation like that in D_3_R, it restricts the flexibility of Trp100^EL1.50^ (Supplementary Movie 3). This trend is consistent with the D_3_R/eticlopride simulations in which we do not observe any significant rearrangement of Trp96^EL1.50^ (Fig. 4d,e,f).

Thus, we infer that the distinct conformation of Trp100^EL1.50^ in the D_2_R structure is a combined effect of the helical EL2 conformation and the favored interaction that Trp100^EL1.50^ can form with bound risperidone in the crystal structure, the latter of which however, has a minimal influence on the binding affinity of risperidone^10^, consistent with the unstable interaction between risperidone and Trp100^EL1.50^ in our simulations (Fig. 4, Supplementary Movie 2). The mutation of this residue to alanine, leucine or phenylalanine did, however, cause substantial increases in both the association and dissociation rate of risperidone^10^. Thus, it appears that the conformation of EL2 influences the dynamics of Trp100^EL1.50^, which in turn controls ligand access and egress to and from the OBS. Both the dissociation and association rates of D_2_R antagonists used as antipsychotics have been proposed to determine their propensity to cause extrapyramidal side-effects and hyperprolactinaemia^24^. Understanding the relationship between the distinct inactive D_2_R conformations stabilized by different antagonist scaffolds and these kinetic parameters will likely be important to facilitate the design of D_2_R antagonists with an optimal kinetic profile that minimizes the risk of side effects.

### EL2 and EL1 dynamics

Next, we evaluated the tendency of the EL2 helix to unwind in each of the simulated D_2_R complexes, by measuring the stability of the backbone H-bond between Ile183^EL2.51^ and Asn186^EL2.54^, a key stabilizing force of the helix (Fig. 5a). When we plotted the Ile183^EL2.51^-Asn186^EL2.54^ distance against the Asp108^3.26^-Ile183^EL2.51^ distance for each D_2_R complex (Fig. 5b), we found the loss of the Asp108^3.26^-Ile183^EL2.51^ interaction increases the probability of breaking the Ile183^EL2.51^-Asn186^EL2.54^ H-bond, i.e., the unwinding of EL2. Interestingly, in all our simulated D_2_R complexes, EL2 has a clear tendency to unwind, regardless of the scaffold of the bound ligand (Fig. 5c,d, Supplementary Movies 1 and 3). Note in the D_3_R/eticlopride simulations, the aligned residues Ser182^EL2.51^ and Asn185^EL2.54^ does not form such a H-bond, and EL2 is always in an extended conformation (Fig. 5b-d). This trend of EL2 to transition towards the extended conformation is also present in our simulations of D_2_R in complex with a partial agonist, aripiprazole, whereas EL2 in the D_3_R complexes with partial agonists (R22 and S22) remains in the extended conformation (Supplementary Table 1 and Supplementary Fig. 5). Interestingly, Asp104^3.26^ and Ser182^EL2.51^ can move into interacting range in the D_3_R/eticlopride simulations, and the Ser182^EL2.51^-Asn185^EL2.54^ interaction can sporadically form in the D_3_R/R22 simulations – both raise a possibility that the extended conformation of D_3_R EL2 may transition to a helical conformation. In addition, while differences in the probabilities of unwinding for each D_2_R complex in Fig. 5 were not drastic, this may be a result of insufficient sampling and cannot exclude the possibility that different ligands may favor particular conformational equilibrium of EL2.

**Figure 5.**
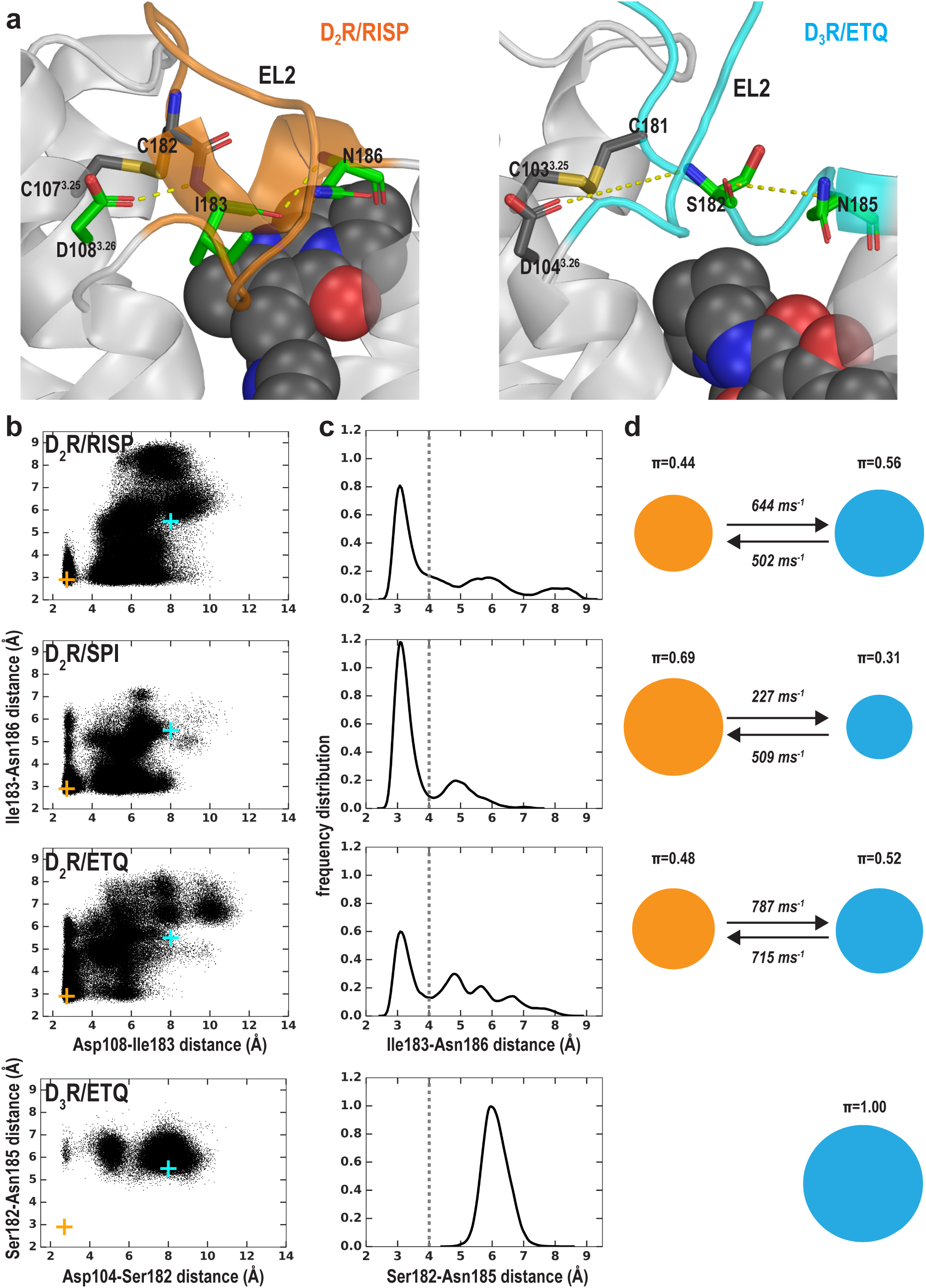
The helical conformation of EL2 in the D_2_R/risperidone structure has a tendency to unwind in our simulations, regardless of the bound ligand. (**a**) The Ile183^EL2.51^-Asn186^EL2.54^ backbone H-bond and the Ile183^EL2.51^-Asp108^3.26^ interaction in D_2_R and their aligned interactions in D_3_R. (**b**) the scatter plots of the two distances in the indicated D_2_R and D_3_R complexes. The orange and cyan crosses indicated the distances in the D_2_R/risperidone and D_3_R/eticlopride structures, respectively. (**c**) The distributions of the EL2.51-EL2.54 distances in the indicated simulations. These distances were used to evaluate the tendency to unwind using Markov state model (MSM) analysis in **d**. (**d**) The MSM analysis of the transition between the helical and extended conformational states of EL2. The area of each disk representing a state is proportional to the equilibrium probability (π) in each simulated condition. The values from the maximum likelihood Bayesian Markov model for π and transition rates from 500 Bayesian Markov model samples are shown. Thus, EL2 in all the D_2_R complexes show significant tendencies to unwind, while that in D_3_R/eticlopride remains extended.

In the fully extended EL2 conformation in which Ile183^EL2.51^ rotates to face the extracellular vestibule, Ile183^EL2.51^ makes a direct contact with the bound risperidone, whereas Trp100^EL1.50^ loses its interaction with the ligand entirely (Supplementary Movie 1). Nevertheless, risperidone retains all other contacts in the OBS. In the recently reported 5-HT_2A_R/risperidone structure (PDB: 6A93)^20^, risperidone has a very similar pose in the OBS as that in the D_2_R structure, occupying the Ile^3.40^ sub-pocket as well. However, on the extracellular side of the OBS, EL2 in the 5-HT_2A_R/risperidone complex is in an extended conformation and the EL2 residue Leu228^EL2.51^ contacting risperidone aligns to Ile183^EL2.51^ of D_2_R, whereas the conserved Trp141^EL1.50^ does not interact with risperidone in the 5-HT_2A_R. It is tempting to speculate the EL2 and EL1 dynamics we observe in the D_2_R/risperidone simulations represents a more comprehensive picture, as the divergent interactions shown in the extracellular loops of the 5-HT_2A_R/risperidone and D_2_R/risperidone structures may not result from differences in recognizing the ligand but rather two different static snapshots due a variety of differences in the crystallographic conditions (Note risperidone has similarly high affinities for both D_2_R and 5HT_2A_R^10,20^). Interestingly, in one of our long MD trajectories of the D_2_R/risperidone complex, EL2 evolved into a conformation that has a helical N-terminal portion and an extended C-terminal portion (Supplementary Movie 4 and Supplementary Fig. 6). This conformation is not observed in either of the D_2_R/risperidone and D_3_R/eticlopride structures but is similar to that of the 5-HT_2A_R/risperidone structure, further demonstrating the dynamics of this loop region (Supplementary Fig. 6).

Thus, the plasticity of the OBS and the dynamics of the extracellular loops appear to be two associated modules in ligand recognition. To the extent of our simulations, we did not detect strong ligand-dependent bias in the EL2 dynamics as we did for the OBS. However, when EL2 is helical, the EL1 dynamics are sensitive to the bound ligand (compare Fig. 4 and Supplementary Fig. 4); when EL2 is extended, it restricts the EL1 dynamics (Fig. 4).

## DISCUSSION

Our results highlight unappreciated conformational complexity for the inactive state of GPCRs and suggest that the risperidone bound D_2_R structure represents only one of a number of possible inactive conformations. Critically, this conformation is incompatible with the binding of other high affinity D_2_R ligands such as eticlopride. While distinct conformational states responsible for functional selectivity have garnered great attention, the potential existence of divergent inactive conformations is of critical importance for high-throughput virtual screening campaigns, as important hits (such as eticlopride) would be missed by focusing on a single antagonist bound state captured in a crystal structure with risperidone, an antagonist with a different scaffold. Such conformations may also confer differences in binding on and off rate or inverse agonist efficacy^25^, that in turn may confer distinct physiological or therapeutic effects. Furthermore, rational lead optimization requires rigorous physical description of molecular recognition^26^, which depends on adequate understanding of the conformational boundary and flexibility of the targeted state.

Previously, using the substituted-cysteine accessibility method (SCAM) in D_2_R ^27,28^, we found that G173^EL2.37^C, N175 ^EL2.39^C, and I184^EL2.52^C were accessible to the MTS reagents and that this accessibility could be blocked by the bound antagonist sulpiride, consistent with their water accessibility and involvement in ligand binding and not with a static orientation facing lipid, whereas A177^EL2.45^C and I183^EL2.51^C were accessible but not protected by sulpiride. Curiously, in the D_2_R/risperidone structure, Ile184^EL2.52^ is only marginally in contact with the ligand, Ile183^EL2.51^ blocks the accessibility of Gly173^EL2.37^ to the OBS and is itself buried in a hydrophobic pocket, whereas Asn175^EL2.49^ faces lipid, where it would be much less reactive. In the D_3_R/eticlopride structure, Ile183^EL2.52^ is in close contact with the bound ligand, Ser182^EL2.51^ faces the extracellular vestibule, whereas the sidechain of Asn173^EL2.39^ is oriented towards the OBS. Thus, it appears that the accessibility pattern of EL2 revealed by previous SCAM studies in D_2_R are more consistent with the extended EL2 conformation revealed by the D_3_R/eticlopride but not with the static D_2_R/risperidone structure (Supplementary Fig. 7). Indeed, we observed spontaneous transitions from the C-terminal helical EL2 conformation to a C-terminal extended conformation in our D_2_R simulations, which suggests that the EL2 conformation of D_2_R exists in an ensemble of structured and unwound conformations, with substantial occupation of the configuration found in D_3_R. Such dynamics of EL2 would explain why the binding of non-selective ligands, such as risperidone or eticlopride, can result in drastically different conformations between the D_2_R and D_3_R structures near EL2, which are not related to the divergence of the receptors. Thus, the D_2_R EL2 appears to have quite dramatic dynamics that are not captured by the crystal structure.

In marked contrast to the obvious trend toward unwinding of EL2 in all our simulated D_2_R complexes, in our recent simulations of MhsT, a transporter protein with a region found by crystallography to alternate between helical and unwound conformations^29^, we failed to observe any spontaneous unwinding over a similar simulation timescale (with the longest simulations being ~5-6 μs) when the region was started from the helical conformation^30,31^. This suggests that the C-terminal helical conformation of EL2 represents a higher energy state than the extended conformation, which allows for observation of the transitions in a simulation timescale not usually adequate to sample folding/unfolding events^32^.

Taken together, our findings reveal that both the plasticity of the transmembrane domain in accommodating different scaffolds and the dynamics of EL2 and EL1 are important considerations in RDD targeting the inactive conformation of D_2_R. More extensive simulations and additional crystal structures bound with ligands from different scaffolds will help to fully reveal the correlations between these two modules.

## METHODS

### Residue indices in EL1 and EL2

Based on a systematic analysis of aminergic receptors, we found a Trp in the middle of EL1 and the disulfide-bonded Cys in the middle of EL2 are the most conserved residues in each segment, and defined their residue indices as EL1.50 and EL2.50, respectively^5^, In this study, for the convenience of comparisons among D_2_R, D_3_R, and D_4_R, and 5-HT_2A_R, based on the alignments of EL1 And EL2 shown in Supplementary Fig. 3, we index the EL1 and EL2 residues of each receptor in the same way as the Ballesteros-Weinstein numbering, e.g., the residues before and after the EL2.50 are EL2.49 and EL2.51, respectively. Note the indices for the shorter sequences are not be consecutive, given the gaps in the alignment.

### Molecular modeling and docking

The D_2_R models in this study are based on the corrected crystal structure of D_2_R bound to risperidone (PDB: 6CM4)^10^. We omitted T4 Lysozyme fused into intracellular loop 3. Three thermostabilizing mutations (Ile122^3.40^A, L375^6.37^A, and L379^6.41^A) were reverted to their WT residues. The missing N terminus in the crystal structure was built de novo using Rosetta^33^, and then integrated with the rest of the D_2_R model using Modeller^34^. Using Modeller, we also extended two helical turns at the TM5 C terminus and threes residues at the TM6 N terminus of the structure and connected these two ends with a 9 Gly loop, similar to our experimentally validated treatment of D_3_R models^35^. The position of the Na^+^ bound in the canonical Na^+^ binding site near the negatively charged Asp^2.50^ was acquired by superimposing the Na^+^ bound structure of adenosine A_2A_ receptor^36^ to our D_2_R models.

The binding poses of risperidone and eticlopride were taken according to their poses in the D_2_R^10^ and D_3_R^9^ structures, respectively. Docking of spiperone in our D_2_R model was performed using the induced-fit docking (IFD) protocol^37^ in the Schrodinger software (release 2017-2; Schrodinger, LLC: New York NY). Based on our hypothesis regarding the role of the Ile^3.40^ sub-pocket in the Na^+^ sensitivity (see text), from the resulting poses of IFD, we choose the spiperone pose with the F-substitution on the butyrophenone ring occupying the Ile^3.40^ sub-pocket. Note that in risperidone and spiperone the F-substitutions have similar distances to the protonated N atoms that interact with Asp^3.32^ (measured by the number of carbon atoms between them, Supplementary Fig. 1).

### Molecular dynamics (MD) simulations

MD simulations of the D_2_R and D_3_R complexes were performed in the explicit water and 1-palmitoyl-2-oleoylphosphatidylcholine (POPC) lipid bilayer environment using Desmond MD System (version 4.5; D. E. Shaw Research, New York, NY) with either the OPLS3e force field^38^ or the CHARMM36 force field^39–42^ and TIP3P water model. For CHARMM36 runs, the eticlopride parameters were obtained through the GAAMP server^43^, with the initial force field based on CGenFF assigned by ParamChem^44^. The system charges were neutralized, and 150 mM NaCl was added. Each system was first minimized and then equilibrated with restraints on the ligand heavy atoms and protein backbone atoms, followed by production runs in an isothermal–isobaric (NPT) ensemble at 310 K and 1 atom with all atoms unrestrained, as described previously ^19,35^. We used Langevin constant pressure and temperature dynamical system ^45^ to maintain the pressure and the temperature, on an anisotropic flexible periodic cell with a constant-ratio constraint applied on the lipid bilayer in the X-Y plane. For each condition, we collected multiple trajectories, the aggregated simulation length is ~340 μs (Table S1).

While the majority of our D_2_R simulations in this study used the OPLS3 force field, to compare with the D_3_R simulations using CHARMM36 that have been continued from the previously reported shorter trajectories^19,35^, we carried out the D_2_R/eticlopride simulations using both the OPLS3 and CHARMM36 force fields (see Table S1). We did not observe significant differences and pooled their results together for the analysis.

### Conformational analysis

Distances and dihedral angles of MD simulation results were calculated with MDTraj (version 1.8.2)^46^ in combination with *in-house* Python scripts.

To characterize the structural changes in the receptor upon ligand binding, we quantified differences of structural elements between the D_2_R/eticlopride and D_2_R/risperidone conditions (using last 600 ns from a representative trajectory for each condition), by applying the previously described pairwise interaction analyzer for GPCR (PIA-GPCR)^35^. The subsegments on the extracellular side of D_2_R were defined as following: TM1e (the extracellular subsegment (e) of TM1, residues 31-38), TM2e (residues 92-96), TM3e (residues 104-113), TM4e (residues 166-172), TM5e (residues 187-195), TM6e (residues 364-369), and TM7e (residues 376-382).

For the PIA-GPCR analysis and the distance analysis in Fig. 4, we used the set of ligand binding residues previously identified by our systematic analysis of GPCR structures. Specifically, for D_2_R, they are residues 91, 94, 95, 100, 110, 111, 114, 115, 118, 119, 122, 167, 184, 189, 190, 193, 194, 197, 198, 353, 357, 360, 361, 364, 365, 367, 368, 376, 379, 380, 383, 384, 386, and 387; for D_3_R, they are residues 86, 89, 90, 96, 106, 107, 110, 111, 114, 115, 118, 165, 183, 188, 189, 192, 193, 196, 197, 338, 342, 345, 346, 349, 350, 352, 353, 362, 365, 366, 369, 370, 372, and 373.

### Markov State Model (MSM) analysis

The MSM analysis was performed using the pyEMMA program (version 2.5.5)^47^. To characterize the dynamics of EL2 of D_2_R, specifically the transitions between helical and extended conformations of its C-terminal portion, we focused on a key hydrogen bond formed in the helical conformation between the backbone carbonyl group of Ile183 and the backbone amine group of Asn186. Thus, for each of the simulated conditions, the distance of Ile183-Asn186 (Ser182-Asn185 in D_3_R) was used as an input feature for the MSM analysis. We discretized this feature into two clusters – distances below and above 4 Å (i.e. EL2 forming a helical conformation and unwinding). Implied relaxation timescale (ITS)^48^ for the transition between these clusters was obtained as a function of various lag times. Convergences of ITS for the MSMs for all conditions was achieved at a lag time of 300 ns (Supplementary Fig. 8), which we further used to estimate Bayesian Markov models with 500 transition matrix samples^49^. The maximum likelihood transition matrix was used to calculate the transition and equilibrium probabilities (π) shown in Fig. 5 and Supplementary Fig. 5.

### [^3^H]spiperone binding assay

For saturating binding assays cell membranes (SNAP-D_2s_-FlpIn CHO, 2.5 μg) were incubated with varying concentrations of [^3^H]spiperone and 10 μM haloperidol as a non-specific control, in binding buffer (20 mM HEPES, 100 mM NaCl, 6 mM MgCl_2_, 1mM EGTA, and 1mM EDTA, pH 7.4) to a final volume of 200 μL and were incubated at 37 °C for 3 h. For competition binding assays cell membranes (SNAP-D_2s_-FlpIn CHO, 2.5 μg) were incubated with varying concentrations of test compound in binding buffer containing 0.2 nM of [^3^H]spiperone to a final volume of 200 μL and were incubated at 37 °C for 3 h. Binding was terminated by fast-flow filtration using a Uniplate 96-well harvester (PerkinElmer) followed by five washes with ice-cold 0.9% NaCl. Bound radioactivity was measured in a MicroBeta2 LumiJET MicroBeta counter (PerkinElmer).

### Data analysis

The concentration of ligand that inhibited half of the [^3^H]spiperone binding (IC_50_) was determined using the following equation:

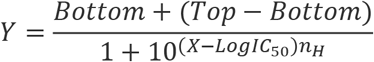

Where Y denotes the percentage specific binding, Top and Bottom denote the maximal and minimal asymptotes, respectively, IC_50_ denotes the X-value when the response is midway between Bottom and Top, and *n*H denotes the Hill slope factor. IC_50_ values obtained from the inhibition curves were converted to *K*_i_ values using the Cheng and Prusoff equation.

## Acknowledgements

Support for this research was provided by the National Institute on Drug Abuse–Intramural Research Program, Z1A DA000606-03 (L.S.), NIH grant MH54137 (J.A.J.) and National Health and Medical Research Council (NHMRC) Project Grant APP1049564 (J.R.L)

## Author Contributions

J.R.L., J.A.J., and L.S. designed the studies. A.M.A. R.K.V., and L.S. performed computational modeling, simulations, and analysis. J.R.L. and H.D.L. performed the binding assay. J.R.L., J.A.J., and L.S. wrote the manuscript, with contributions from all the authors. Jackie Glenn is thanked for technical support in generating membrane preparations.

## SUPPLEMENTARY INFORMATION

**Supplementary Table 1.** Summary of molecular dynamics simulations.

| receptor | ligand | bound<br>Na<sup>+</sup> | number of<br>OPLS3<br>trajectories | Number of<br>CHARMM36<br>trajectories | accumulated<br>simulation time<br>(ns) |
| --- | --- | --- | --- | --- | --- |
| D<sub>2</sub>R | risperidone | + | 12 |  | 29610 |
|  |  | - | 11 |  | 36000 |
|  | spiperone | + | 21 |  | 39900 |
|  |  | - | 17 |  | 25470 |
|  | eticlopride | + | 5 | 12 | 46020 |
|  |  | - | 7 |  | 8400 |
|  | aripiprazole | + | 40 |  | 66660 |
| D<sub>3</sub>R | eticlopride | + |  | 3 | 13200 |
|  |  | - |  | 4 | 6240 |
|  | R22 | + |  | 7 | 33600 |
|  | S22 | - |  | 6 | 49920 |
| Total |  |  | 113 | 32 | 355020 |

**Supplementary Figure 1.**
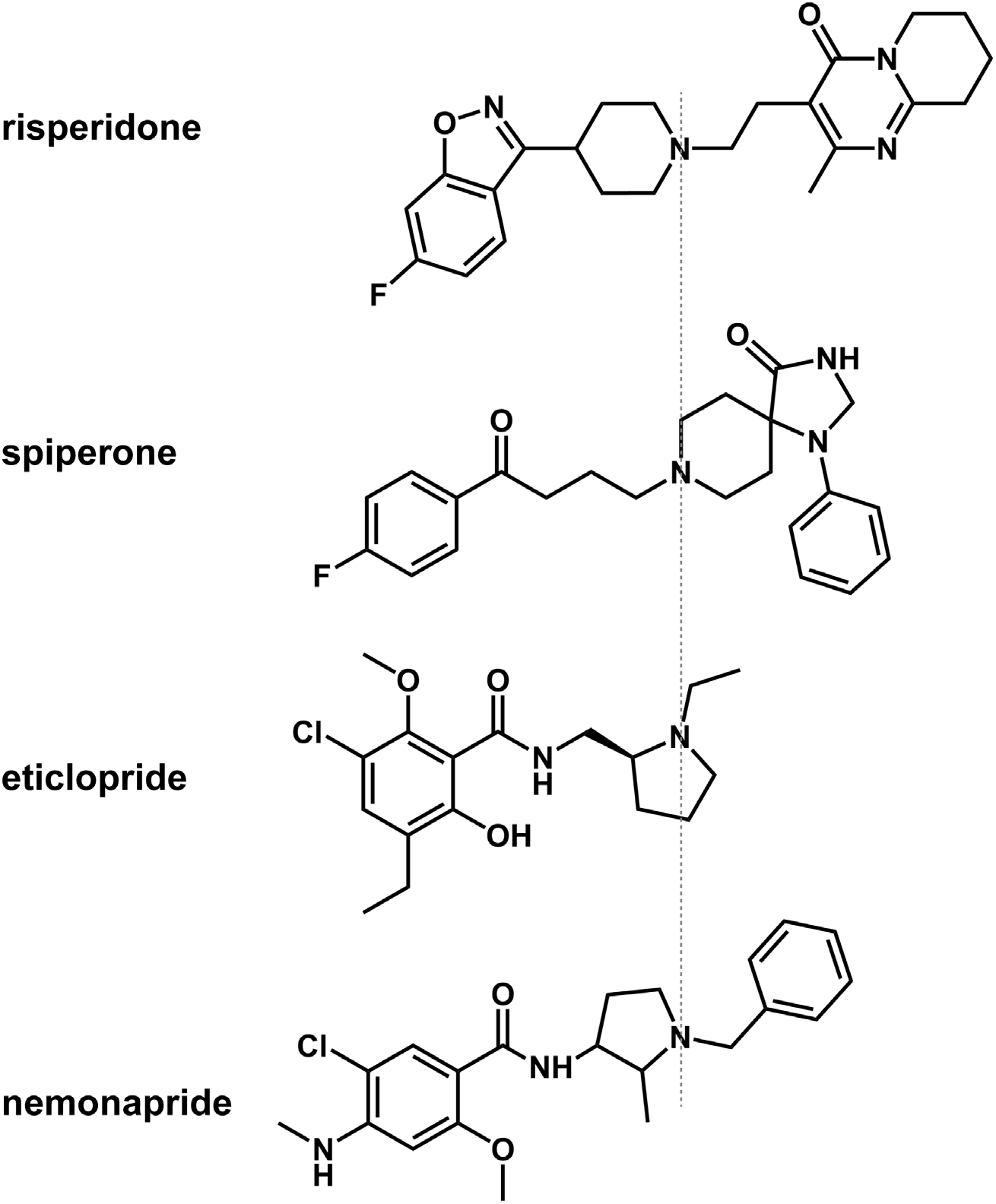
Chemical structure alignments of the non-selective D_2_-like receptors ligands.

**Supplementary Figure 2.**
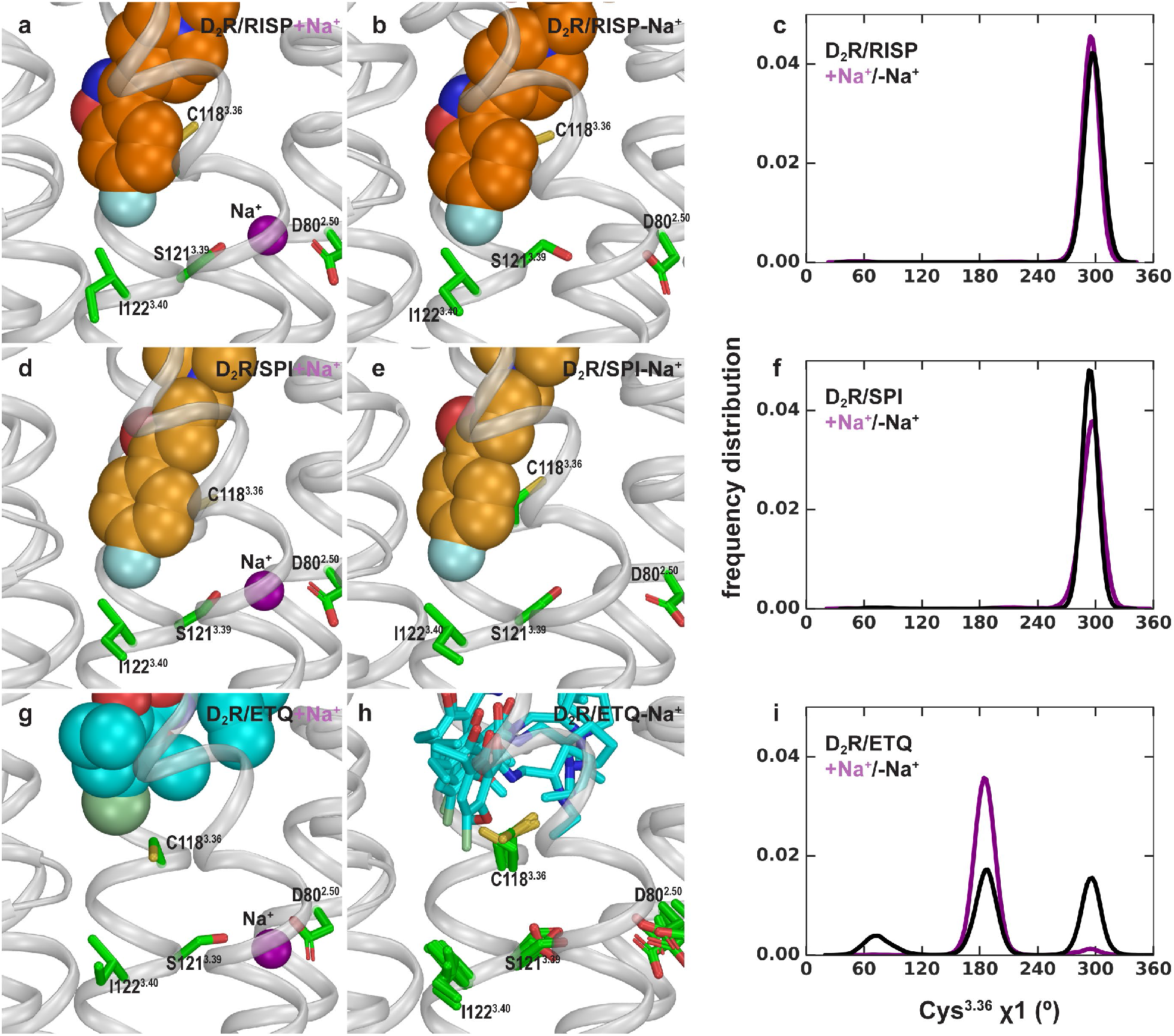
Allosteric communication between the Ile^3.40^ sub-pocket and the Na^+^ binding site. Risperidone (**a**, **b**) and spiperone (**d**, **e**) similarly occupy the Ile^3.40^ sub-pocket in both the presence and absence of Na^+^ bound at the Asp80^2.50^ site. In the eticlopride bound conditions (**g**, **h**), the Ile^3.40^ sub-pocket is not occupied, and Cys^3.36^ shows flexibility in the absence of bound Na^+^. (**c**, **f**, and **i**) Distributions of the χ1 rotamer of Cys^3.36^ in the D_2_R simulations in the presence of different bound ligands.

**Supplementary Figure 3.**
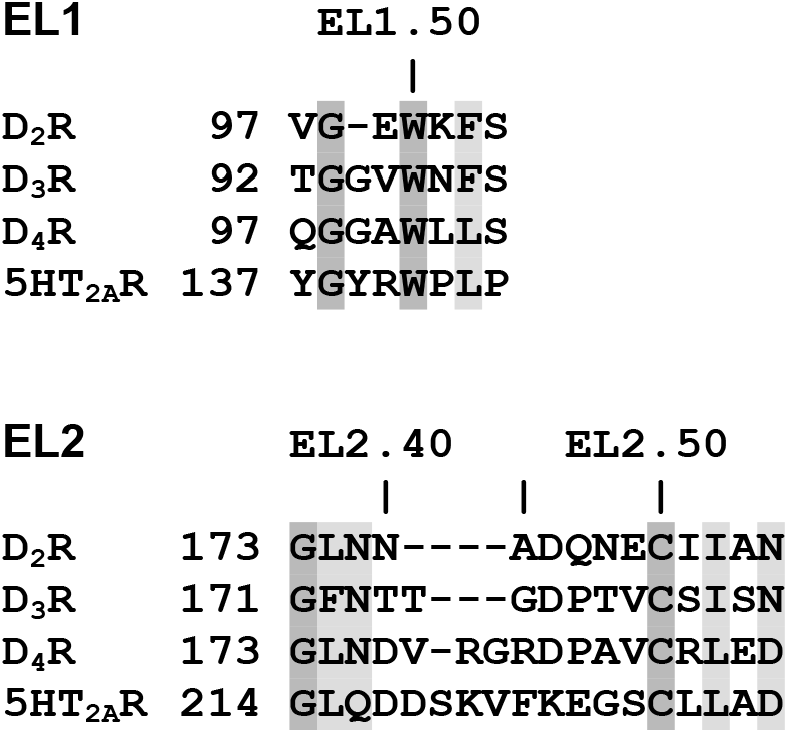
Sequence alignment and residue indices of EL1 and EL2 for the receptors being compared in this study. The positions with identical residues are in dark grey shade, the conserved positions are in light grey shade.

**Supplementary Figure 4.**
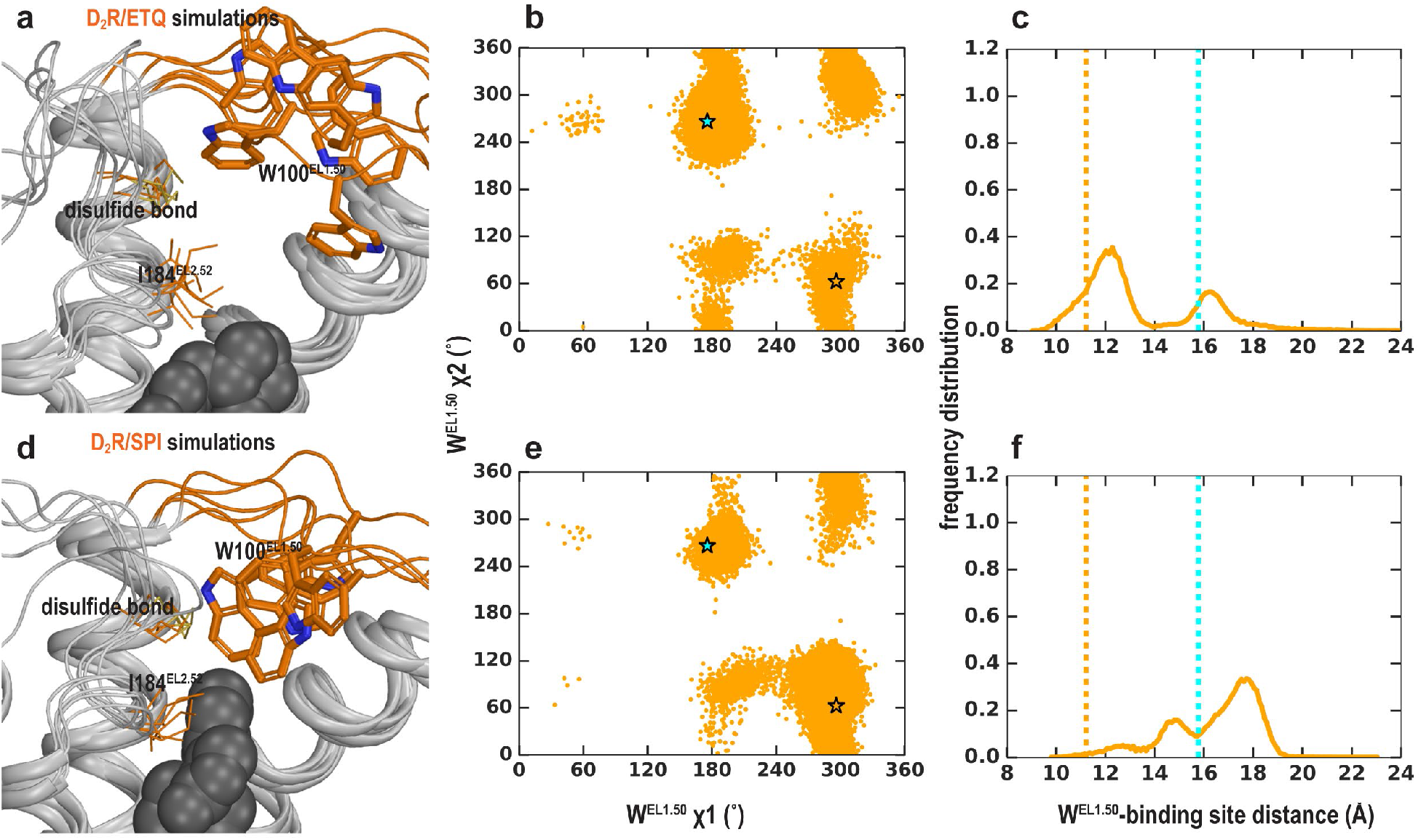
EL1 is dynamic in the D_2_R/eticlopride and D_2_R/spiperone simulations when EL2 is helical. Trp100 shows significant flexibility and can adopt multiple positions and orientations in D_2_R/eticlopride (**a**-**c**) and D_2_R/spiperone (**d**-**f**) simulations. Their χ1 and χ2 dihedral angles of Trp100 (**b**, **e**) and the distance between Trp100 and the ligand binding site (**c**, **f**) have wide and different distributions. These dihedral angle values in the D_2_R and D_3_R structures are indicated with the orange and cyan stars, respectively. The distances in the D_2_R and D_3_R structures are indicated with the orange and cyan dotted lines, respectively.

**Supplementary Figure 5.**
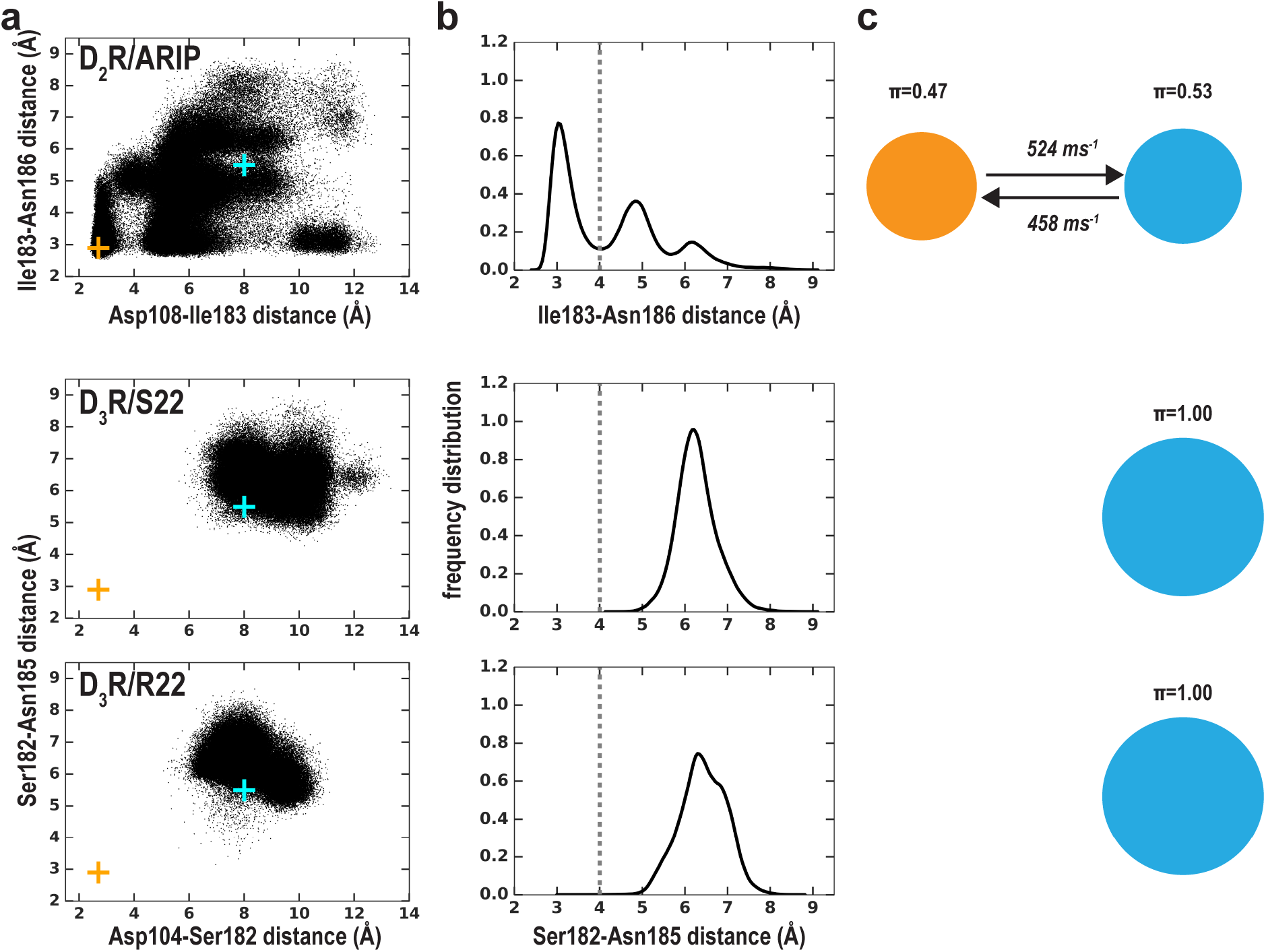
The MSM analysis of Ile183-Asn186 distance in the simulations of the D_2_R/aripiprazole, D_3_R/S22, and D_3_R/R22 complexes (Supplementary Table 1). The early stage of D_3_R/S22 and D_3_R/R22 simulations has been reported previously^35^. The representation and color scheme is the same as that for **Fig. 5**.

**Supplementary Figure 6.**
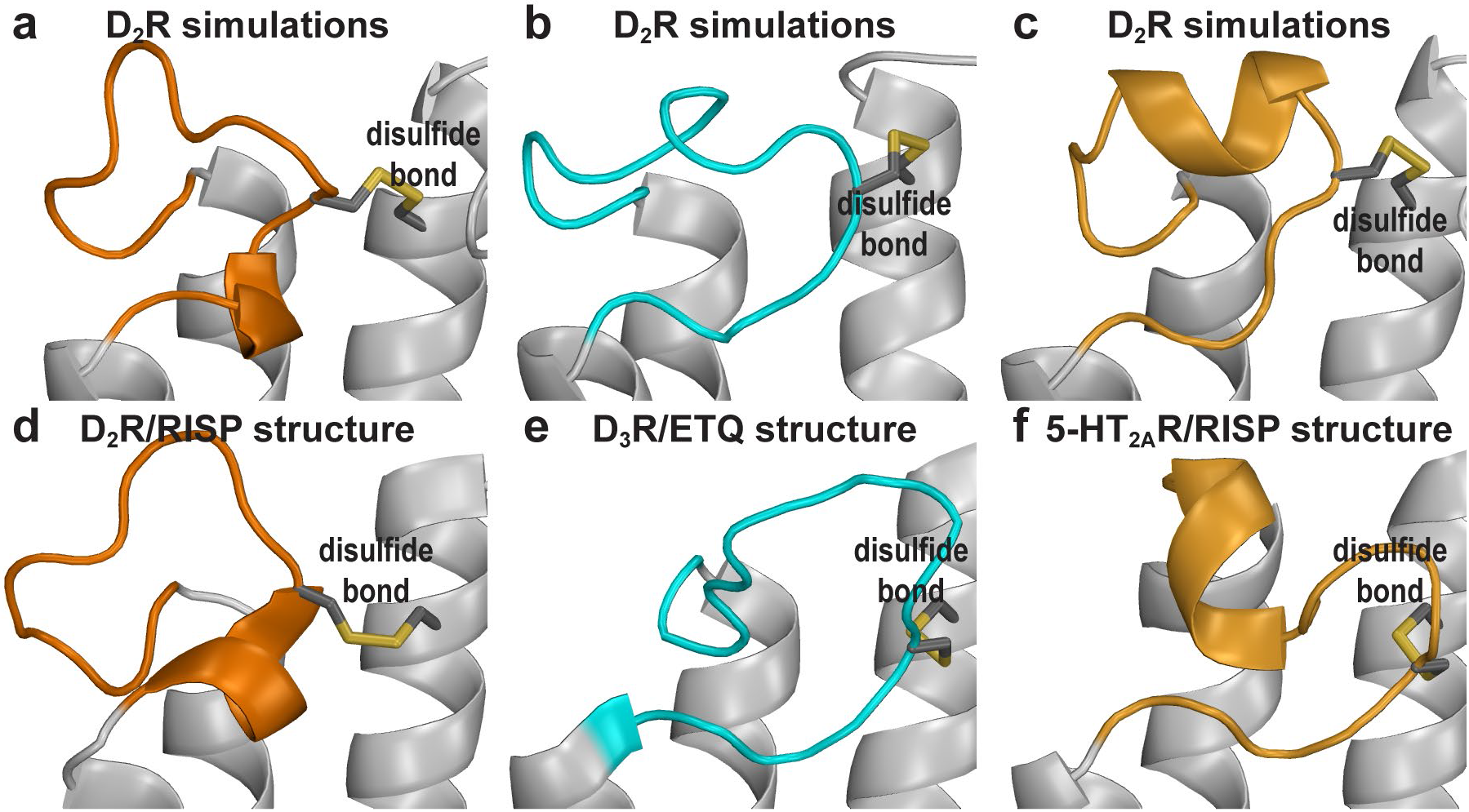
The distinct D_2_R EL2 conformations revealed by the MD simulations are similar to those of homologous receptors. The C-terminal helical EL2 conformation in the D_2_R structure (**d**) can be maintained in the simulations (**a**). the C-terminal extended conformation (**b**) is similar to those in the D_3_R structure (**e**). The N-terminal helical conformation (**c**) is reminiscent of that in the 5-HT_2A_R/risperidone structure (**g**), and those in β_1_ and β_2_ adrenergic receptors structures (not shown).

**Supplementary Figure 7.**
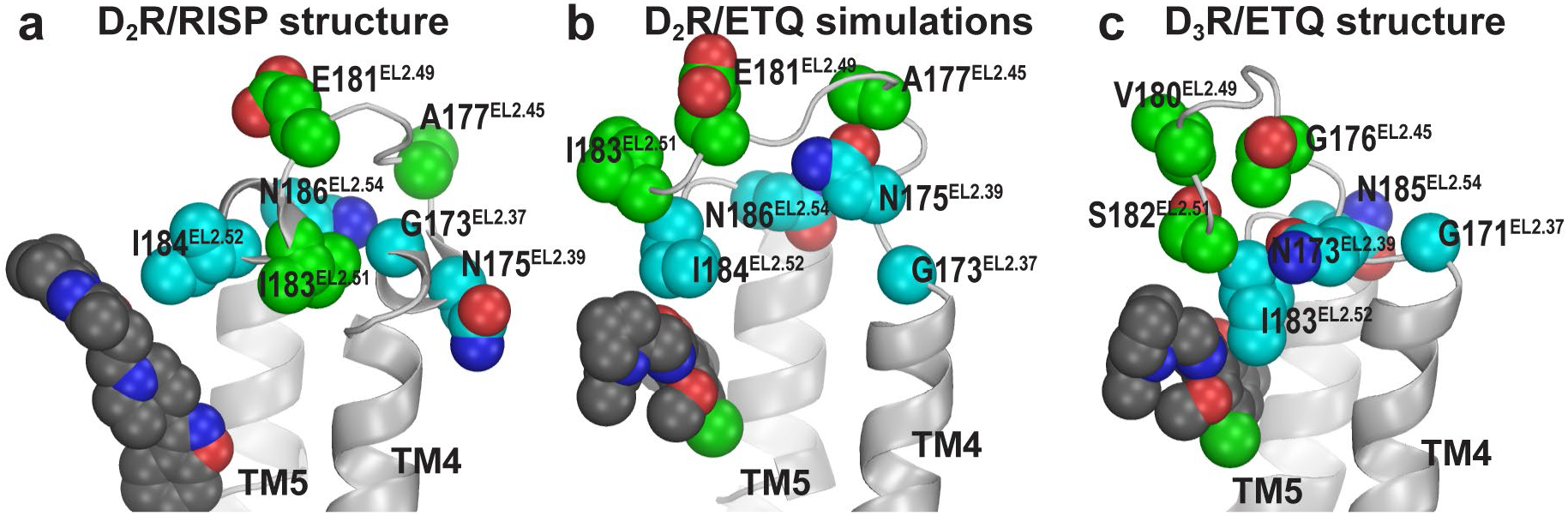
The accessibility pattern of EL2 revealed by previous SCAM studies in D_2_R is more consistent with an extended EL2 conformation similar to that in the D_3_R/eticlopride structure. The accessible residues are in green, the protected residues are in cyan. In the D_2_R/risperidone structure (**a**), Ile183^EL2.51^ blocks the accessibility of Gly173^EL2.37^ to the OBS, while Asn175 faces lipid. In the D_2_R/eticlopride simulations (**b**) and D_3_R/eticlopride structure (**c**), Asn^EL2.39^ rotates to point inward, while Ile183^EL2.51^ in D_2_R and Ser182^EL2.51^ in D_3_R rotates to face the extracellular vestibule of the receptors.

**Supplementary Figure 8.**
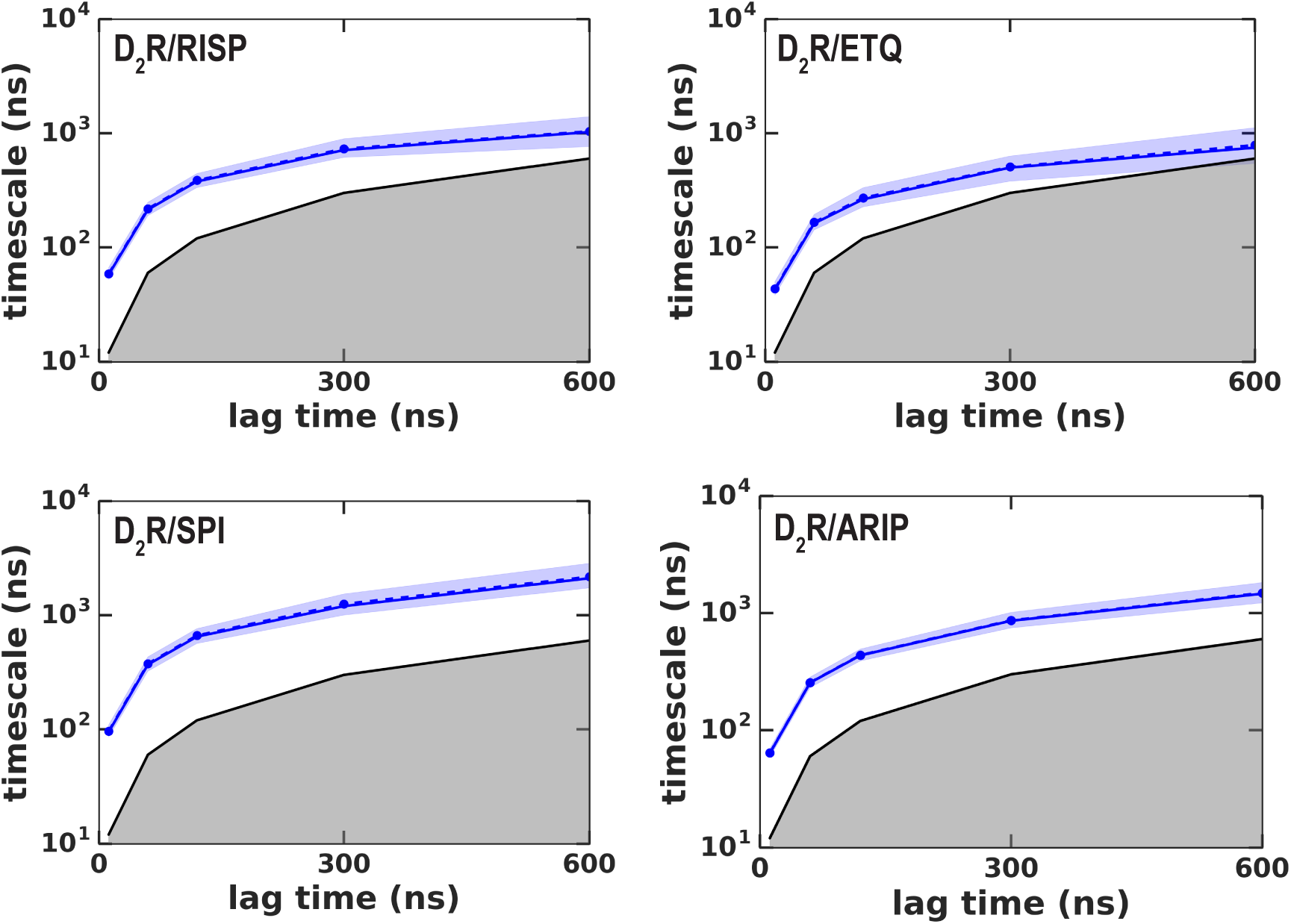
Implied timescales (ITS) for the MSM analysis. The implied timescales (ITS) of the transition between the two states in each of the D_2_R conditions shown in Fig. 5 and Supplementary Fig. 5 are plotted against various lag times. ITSs were not computed for D_3_R conditions because there was not transition between two states. The ITS of the maximum likelihood Bayesian Markov model is shown in a blue solid line, whereas the means and the 95% confidence intervals (computed by Bayesian sampling) are shown in dashed and shaded areas, respectively. Timescales smaller than the lag time are shown in grey-shaded area. A lag time of 300 ns was chosen for our analysis.

**Supplementary Movie 1.** A movie of a 4.2 μs D_2_R/risperidone trajectory collected using the OPLS3 force field shows spontaneous unwinding of EL2. The conformation of EL2 gradually transitions to an extended configuration similar to that in the D_3_R structure. See Fig. 3 for the pathway of unwinding. Note that the extended conformation of EL2 stabilizes Trp100^EL1.50^. The Cα atom of Gly173^EL2.37^, the sidechains of Trp100^EL1.50^, Ile183^EL2.51^, and Ile184^EL2.52^ and the bound risperidone are shown as spheres. Asp108^3.26^ and the disulfide bond between Cys107^3.25^ and Cys182^EL2.50^ are shown as sticks. The carbon atoms of Gly173^EL2.37^ and Ile184^EL2.52^ are colored in cyan, those of Ile183^EL2.51^ are in green, those of Trp100^EL1.50^, Cys107^3.25^, Asp108^3.26^, Asn175^EL2.39^, and Cys182 ^EL2.50^ are in dark grey; those of the bound ligand risperidone are in orange.

**Supplementary Movie 2**. A movie of a 4.2 μs D_2_R/eticlopride trajectory shows the dynamics of Trp100^EL1.50^ when the C-terminal portion of EL2 is in a helical conformation. Note that Trp100^EL1.50^ can be stabilized by interacting with the disulfide bond. The presentation and color scheme are similar to those in Supplementary Movie 1, except that the bound carbon atoms of the ligand eticlopride are colored in cyan.

**Supplementary Movie 3**. A movie of a 3.6 μs D_2_R/eticlopride trajectory collected using the CHARMM36 force field shows another example of unwinding of EL2. Thus, considering the similar unwinding pathway as that in Movie S1 (Fig. 3), the unwinding does not depend on the force field used in the simulations or the identity of the antagonist bound in the OBS. Note the sidechain of Asn175^EL2.39^ rotates inward and approaches Asp108^3.26^ in this trajectory. The presentation and color scheme are the same as those in Supplementary Movie 2.

**Supplementary Movie 4.** A movie of a 4.5 μs D_2_R/risperidone trajectory shows the N-terminal portion of EL2 can transition into a helical conformation when the C-terminal portion is extended. This is a novel EL2 conformation that has not been revealed by the D_2_R, D_3_R or D_4_R structures but similar to those in the 5-HT_2A_R/risperidone, β_2_AR and β_2_AR structures (Supplementary Fig. S6). The presentation and color scheme are the same as those in Supplementary Movie 1.

